# Unique mapping of structural and functional connectivity on cognition

**DOI:** 10.1101/296913

**Authors:** J. Zimmermann, J.G. Griffiths, A.R. McIntosh

## Abstract

The unique mapping of structural and functional brain connectivity (SC, FC) on cognition is currently not well understood. It is not clear whether cognition is mapped via a global connectome pattern or instead is underpinned by several sets of distributed connectivity patterns. Moreover, we also do not know whether the pattern of SC and of FC that underlie cognition are overlapping or distinct. Here, we study the relationship between SC and FC and an array of psychological tasks in 609 subjects from the Human Connectome Project (HCP). We identified several sets of connections that each uniquely map onto different aspects of cognitive function. We found a small number of distributed SC and a larger set of cortico-cortical and cortico-subcortical FC that express this association. Importantly, SC and FC each show unique and distinct patterns of variance across subjects and differential relationships to cognition. The results suggest that a complete understanding of connectome underpinnings of cognition calls for a combination of the two modalities.

**Significance Statement:** Structural connectivity (SC), the physical white-matter inter-regional pathways in the brain, and functional connectivity (FC), the temporal co-activations between activity of brain regions, have each been studied extensively. Little is known, however, about the distribution of variance in connections as they relate to cognition. Here, in a large sample of subjects (N = 609), we showed that two sets of brain-behavioural patterns capture the correlations between SC, and FC with a wide range of cognitive tasks, respectively. These brain-behavioural patterns reveal distinct sets of connections within the SC and the FC network and provide new evidence that SC and FC each provide unique information for cognition.

## Introduction

In neuroscience, big data initiatives such as the Human Connectome Project (HCP) acquire connectomic and phenotypic data from a large number of individuals in an effort to understand how brain networks relate to individual behaviour (Van Essen et al., 2013; Fornito, 2016). Connectomes can represent either structural connectivity (SC), the white-matter inter-regional pathways estimated from diffusion-weighted MR imaging (Baldassarre et al., 2012), or functional connectivity (FC), the patterns of temporal dependencies between regional activity measurements such as blood oxygen level dependent functional magnetic resonance imaging (fMRI) timeseries. The most commonly studied form of FC, resting-state FC (rsFC), is measured in the absence of an explicit task. It represents meaningful coordinated fluctuations that have been related to task FC and performance (Mennes et al., 2010; Baldassarre et al., 2012; Stevens and Spreng, 2014).

Both SC (Matejko et al., 2013; Willmes et al., 2014; Moeller et al., 2015; Klein et al., 2016) and rsFC (Song et al., 2008; Song et al., 2009; Pamplona et al., 2015; Santarnecchi et al., 2015; Hearne et al., 2016; Smith, 2016; Ferguson MA, 2017; Pezoulas et al., 2017) have been linked to cognitive functioning, including higher order cognitive processes such as fluid and crystallized intelligence, visuospatial processing (Ponsoda et al., 2017), or numerical cognition (Matejko et al., 2013; Willmes et al., 2014; Moeller et al., 2015; Klein et al., 2016).

What remains unclear, however, is the specificity of the behaviourally-relevant aspect of the connectome. That is, can individual variability across connectomes be explained by a single set of connections that is highly predictive of cognitive function *more generally*? Or are there multiple sets of connections that *each predict different* aspects of cognitive functioning? The former suggests that connectomes are relevant for understanding general differences in global cognitive functioning whilst the latter suggests that individual differences in particular networks can inform specific types of cognitive functioning, such attention, memory, or executive function. Evidence of both views exists. For instance, Rosenberg et al. (2013) showed that a particular set of connectivity patterns predicted individuals’ attention ability, supporting the notion of specific connectome-behaviour associations. The connections identified were specific to attention ability and did not predict cognition more generally. In contrast, other research has shown that a single mode of covariation can capture relationships between a distributed set of functional connections and a wide set of behavioural and demographic variables (Smith et al., 2015). The same question can be asked for SC.

Currently, little is known about the overlap between the two modalities in their mapping to individual differences in cognition. Given the limitations of mapping individual SC to FC (Koch et al., 2002; Skudlarski et al., 2008; Honey et al., 2009; Zimmermann et al., 2016; Zimmermann et al., 2018), we might expect that the two modalities provide unique and distinct sources of cognitive-related variability (Duda et al., 2010; Hirsiger et al., 2016). On the other hand, cognition arises from an interplay of structure and function, and so a degree of overlap in the spatial pattern of networks that give rise to cognitive function is expected.

In the present study, we examine how cortical and subcortical SC and rsFC from 609 subjects from the Human Connectome Project relates to a wide range of cognitive functions, including working memory, executive function/cognitive flexibility, processing speed, fluid intelligence, episodic memory, and attention/inhibitory control. We examined 1) whether there exists a single set of connections that generally map onto cognition, or rather several sets of connections that map onto different aspects of cognition, and 2) whether the patterns of connectivity that map onto cognition are independent and unique for SC versus rsFC or whether they provide common information. We used Partial Least Squares to map orthogonal patterns of brain-behavioural relationships (McIntosh and Lobaugh, 2004; Krishnan et al., 2011). This method is comparable to canonical correlation, however, is more suitable for neuroimaging data as it is robust to collinearity within the datasets.

## Methods

### Subjects and behavioural measures

The sample included 609 genetically unrelated subjects from the Q7 HCP release (M/F = 269/340 female; age range = 22-36) (Van Essen et al., 2013). The research was performed in compliance with the Code of Ethics of the World Medical Association (Declaration of Helsinki). All subjects provided written informed consent, approved by the ethics committee in accordance with guidelines of HCP WU-Minn. In the current study, 11 behavioural measures of cognitive function were correlated with SC and FC (described below). Cognitive measures covered a range of processes. Note that the working memory 0-back and 2-back tasks each included 2 measures: accuracy and reaction time. The measures are described in Table 1 below and in detail in the following: https://wiki.humanconnectome.org/display/PublicData/HCP+Data+Dictionary+Public+500+Subject+Release. Note that cognitive scores were age adjusted.

**Table 1.**
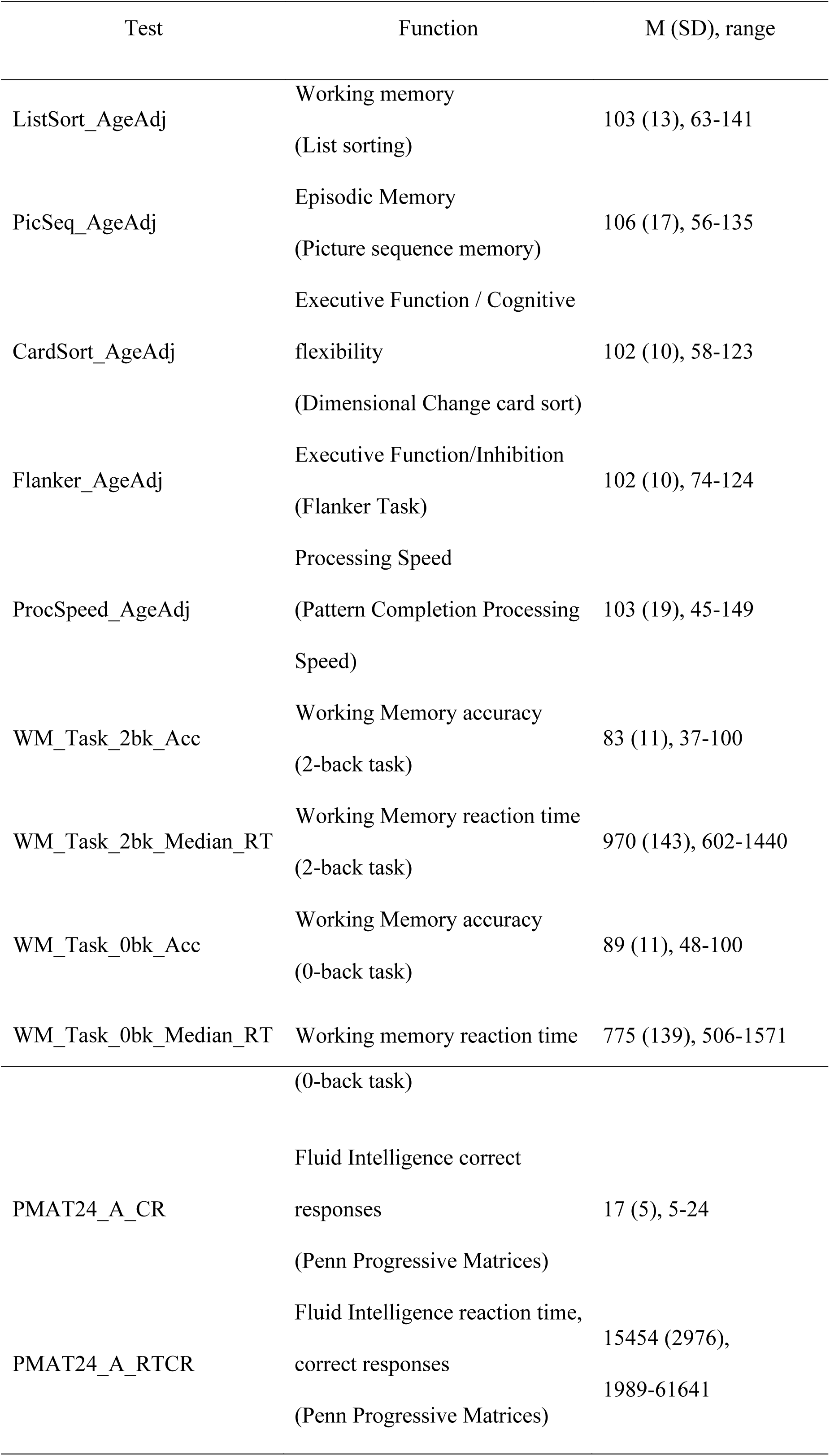
Cognitive tests and corresponding function

### MRI acquisition and preprocessing

Data acquisition details for the WU-Minn HCP corpus (Van Essen et al., 2013) were described in detail elsewhere (Smith et al., 2013; Sotiropoulos et al., 2013; Ugurbil et al., 2013). The present study made use of resting-state BOLD fMRI, multi-shell (multiple non zero b-values) high-resolution dwMRI, and structural (T1-weighted MRI) data from the HCP minimally preprocessed pipeline (Glasser et al., 2013; Smith et al., 2013).

Diffusion data with a spin-echo multiband EPI sequence, with 111 slices of 1.25 mm isotropic voxels (Feinberg et al., 2010; Sotiropoulos et al., 2013; Ugurbil et al., 2013), distortion corrected (Andersson et al., 2003; Andersson and Sotiropoulos, 2015) were used. Segmentation and parcellation was performed on the basis of the high-resolution T1-weighted image (voxel size: 0.7 mm isotropic) of each subject using FreeSurfer (Fischl, 2012), automatically parcellating the brain into 83 cortical and subcortical ROIs (41 per hemisphere, plus brainstem) according to the Lausanne 2008 atlas (Daducci et al., 2012). An 83 × 83 connectivity matrix was formed, representing for each pair of regions their reconstructed pathways. Deterministic tractography using Dipy software (Garyfallidis et al., 2014) was performed on minimally preprocessed data using a method similar to that of Hagmann et al (2008) and Cammoun et al (2012). Streamlines were computed from 60 equally spaced points within each voxel on the grey-white matter interface via the EuDX algorithm (Garyfallidis et al (2012). Streamlines that were shorter than 10mm or longer than 250mm were discarded, as well as those that did not terminate at the grey-white matter interface. The strength of the reconstructed connections was measured as streamline density, computed as the number of tractography streamlines that touched both cortical regions (Hagmann et al., 2008).

Whole brain echo-planar images (EPI) (TR = 720 ms, 2 mm^3^ voxels) (Moeller et al., 2010; Ugurbil et al., 2013) with denoising procedures from the resting-state FIX denoised dataset were used (Glasser et al., 2013; Smith et al., 2013). Data included both phase encoding acquisition directions (L-R, R-L) (acquisition time = 14 mins, 33 sec). We computed average regional timeseries from these voxels, based on the 83 cortical and subcortical regions. Pearson’s correlations were calculated for each pair of timeseries to compute the FC between all regions; these were then Fisher Z-transformed, and the two (L-R and R-L) pairs of matrices were averaged for each subject. The resulting FC matrices were 83×83 for the 609 subjects. Global signal regression was not performed for comparability with previous resting state studies. Note that age was regressed from SC and Fisher’s z-transformed FC, and residuals were used for analysis.

### Experimental Design and Statistical Analysis

#### Cognitive Measures

The 11 cognitive measures were correlated (Pearson’s) with one another to quantify the relationship between them. The resulting p-value cross-correlation matrix was of size 11 × 11. The p-value of each correlation value was corrected for multiple comparisons using FDR (Matlab function fdr_bky) (Benjamini, 2006). We also wanted to understand whether the cognitive measures describe a single global cognitive component of intelligence, or whether numerous components are required to capture the different subtypes of cognitive functioning. To this end, we conducted a principal components analysis (PCA) of the cognitive measures (Matlab function princomp). To assess the significance of the resulting eigenvalues, we permuted the cognitive measures 100 times (scrambled across cognitive measures and subjects) and decomposed (PCA) the permuted cognitive matrices to obtain a null distribution of eigenvalues for each PC. We then calculated the p-value per PC eigenvalue as the proportion of times the permuted eigenvalue exceeded the obtained eigenvalue.

#### Correlation between cognition and connectivity

We used Partial Least Squares (PLS) for neuroimaging (McIntosh and Lobaugh, 2004; Krishnan et al., 2011) in order to assess the multidimensional connectome-cognition relationships. We used the Behavioral PLS correlation function described there, ran in Matlab using custom code. As the connectivity matrices are symmetric, we used the vectorized upper triangle of the connectivity matrices from the 609 subjects. Connections that were 0 for 66% of subjects or more were not included in the analysis, resulting in 870 remaining connections for SC and 3403 connections for FC. Vectorized connectomes were then stacked resulting in a subjects*connections brain matrix, for SC = 609*870, and for FC = 609*3403. The behavioural matrix was subjects*behavioural measures, of size 609*11.

The brain and behavioural matrices were then cross-correlated to start the PLS analysis. PLS is multivariate analysis method comparable to canonical correlation, and captures maximally covarying brain-behaviour relationships in mutually orthogonal latent variables (LVs). The significance of LVs is assessed via permutation tests (1000 iterations) of the singular values from singular value decomposition (SVD) of the brain and behavioural correlation matrices, and reliability of each connectivity estimate to the LV is assessed via bootstrap resampling (3000 iterations). The reliability of each connection’s loading onto the brain-behaviour relationship in each LV is represented as a bootstrap ratio, the ratio of a connection’s weight over its estimated standard error. The ratio can be considered equivalent to a z-score, but is used to impart reliability rather than significance. A connection with a positive high bootstrap ratio contributes positively and reliably to the brain-behaviour correlation obtained for that LV, whereas a connection with a negative high bootstrap ratio contributes negatively and reliably to the brain-behaviour relationship. Bootstrapping is also used to construct confidence intervals on the brain-behaviour correlations.

#### SC-Cognition vs FC-Cognition

In order to compare the connections that contributed to the SC-cognition relationship and those that contributed to the FC-cognition relationship, we calculated the scalar dot product (Matlab function dot) between the brain scores (“U”) from the PLS SVD expressed in the FC-cognition and those expressed in the SC-cognition, across all significant LVs. We did the same thing for cognitive measures between analyses, using Pearson’s correlations instead of dot products. Note that these correlations were corrected for multiple comparisons using FDR (Matlab function fdr_bky) (Benjamini, 2006).

## Results

### Cognitive measures

The cognitive scores correlated positively amongst one another, with the exception of the WM reaction time tasks, which correlated negatively with all other cognitive tests (See Figure 1). All correlations were significant (all *p* < 0.001, MC corrected, See Methods).

**Figure 1.**
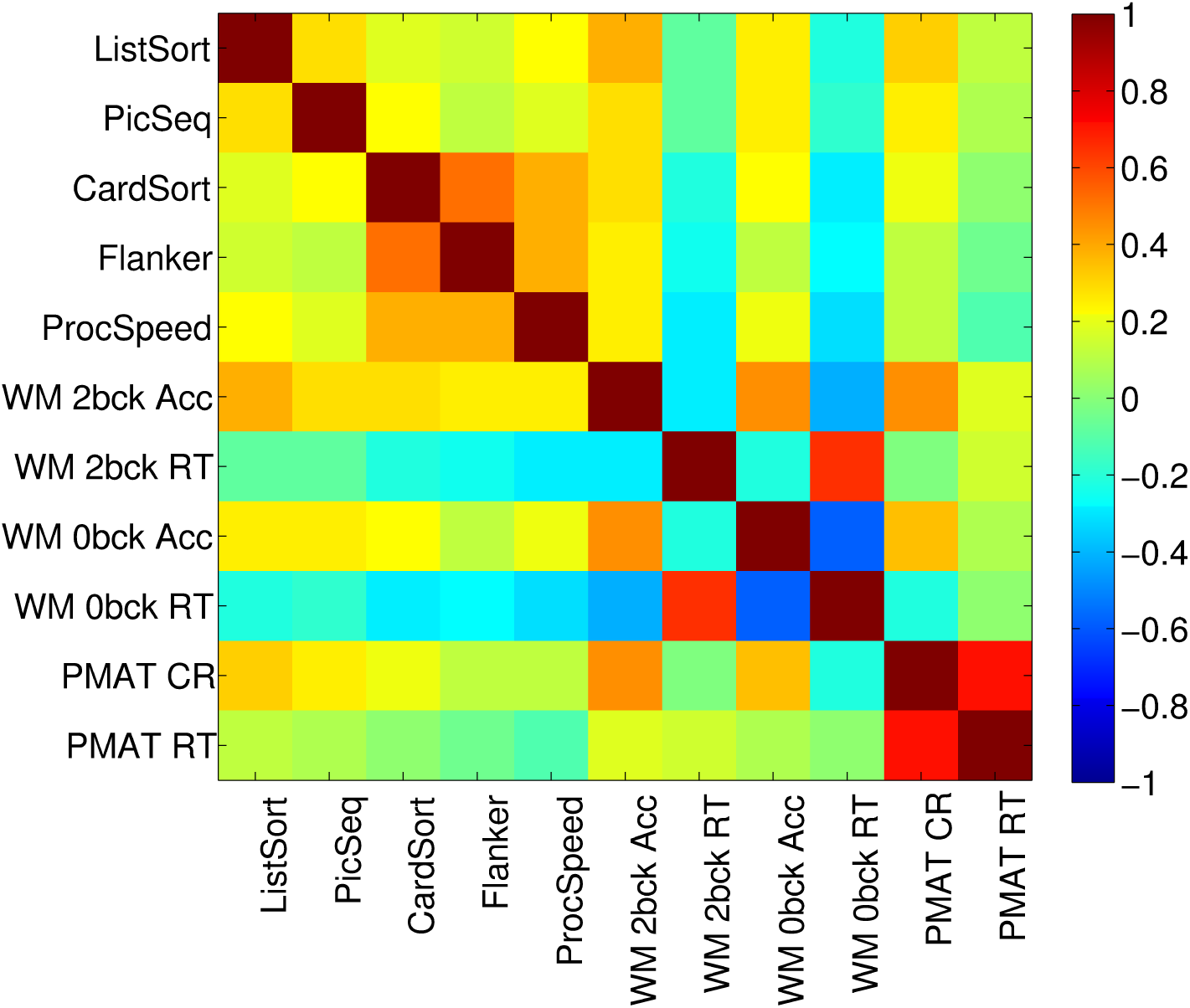
Correlation (Pearson’s) amongst the cognitive tests.

A PCA of the 11 cognitive test scores yielded three significant PCs, where significance was assessed by permutation testing (See Methods) (λ = 3.5072, 1.8679, and 1.2026, % variance explained = 61, 17, 7 respectively) (Figure 2). All cognitive tests loaded onto the first PC, with an emphasis on the speed-accuracy trade-off for the two WM tests. Thus, participants who were more accurate were also slower on these WM tests. The second PC emphasized the PMAT tests; note that reaction time for correct responses (PMAT RT) and the number of correct responses (PMAT CR) were positively correlated. The third PC emphasized similarities between the two WM RT tests and the remaining tests, in opposition to the two WM accuracy measures.

**Figure 2.**
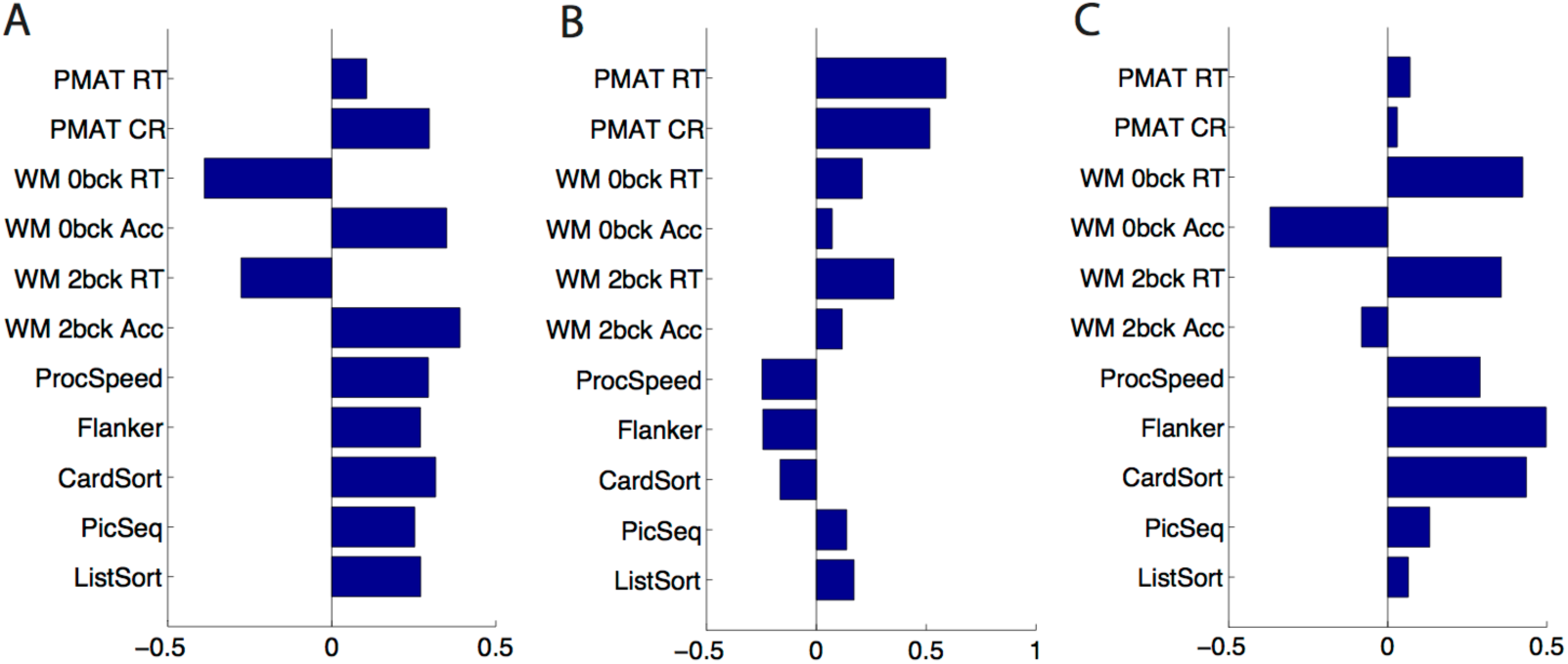
PCA of the 11 cognitive tests. Shown here are the principal component coefficients (loadings) on each PC. A) PC1 B) PC2 C) PC3.

### Correlation between cognition and connectivity

PLS analyses identified two significant LVs that describe the relationship between SC and cognition, and FC and cognition each, respectively.

#### SC-Cognition

The two LVs that captured the SC-cognition association revealed two distinct patterns of cognitive functions that mapped onto two sets of unique structural connections (Figure 3). (LV1 SC: 37.22% of total covariance, singular value = 2.53, *p* = 0.04, LV2 SC: 14.81% of total covariance, singular value = 1.6, *p* = 0.03). For both LVs, a small number of connections across cortical and subcortical regions were stable by bootstrap. Connections loaded positively and negatively onto each of the SC-cognition LVs (See bootstrap ratios in Figure 3B, 3D). For the cognitive measures, the first LV strongly expressed the full array of cognitive tests, with a speed-accuracy trade-off for the WM tests. The second LV expressed primarily the two PMAT tests.

**Figure 3.**
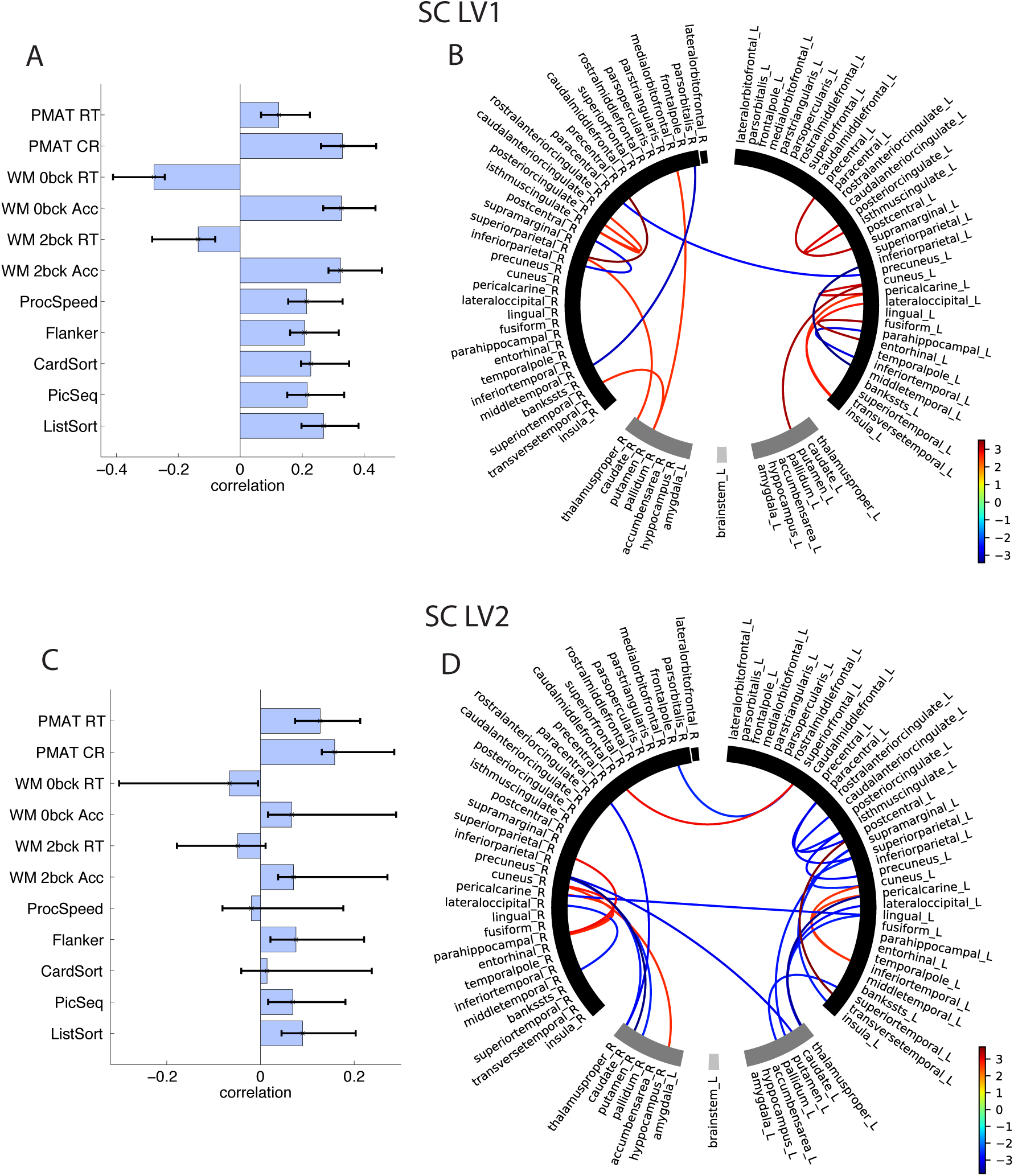
SC-Cognition A) LV1 correlations between SC and cognition, with CIs from bootstrap resampling, B) LV1 bootstrap ratios, these are connection loadings on the SC-cognition relationship, bootstrap threshold = -2.5, 2.5. Connections with positive bootstrap ratios contribute positively to the SC-cognition correlation, negative bootstrap ratios contribute negatively to the SC-cognition correlation. Right and left hemisphere regions, and subcortical and cortical regions, are separated by a space in the circular plot. C) LV2 correlations between SC and cognition D) LV2 bootstrap ratios

#### FC-Cognition

The two LVs that captured the FC-cognition association revealed two distinct patterns of cognitive functions that map onto two sets of functional connections (Figure 4). (LV1 FC: 41.5% of total covariance, singular value = 6.45, *p* = 0.01 LV2 FC: 33% of total covariance, singular value = 5.75, *p* = 0.046).

**Figure 4.**
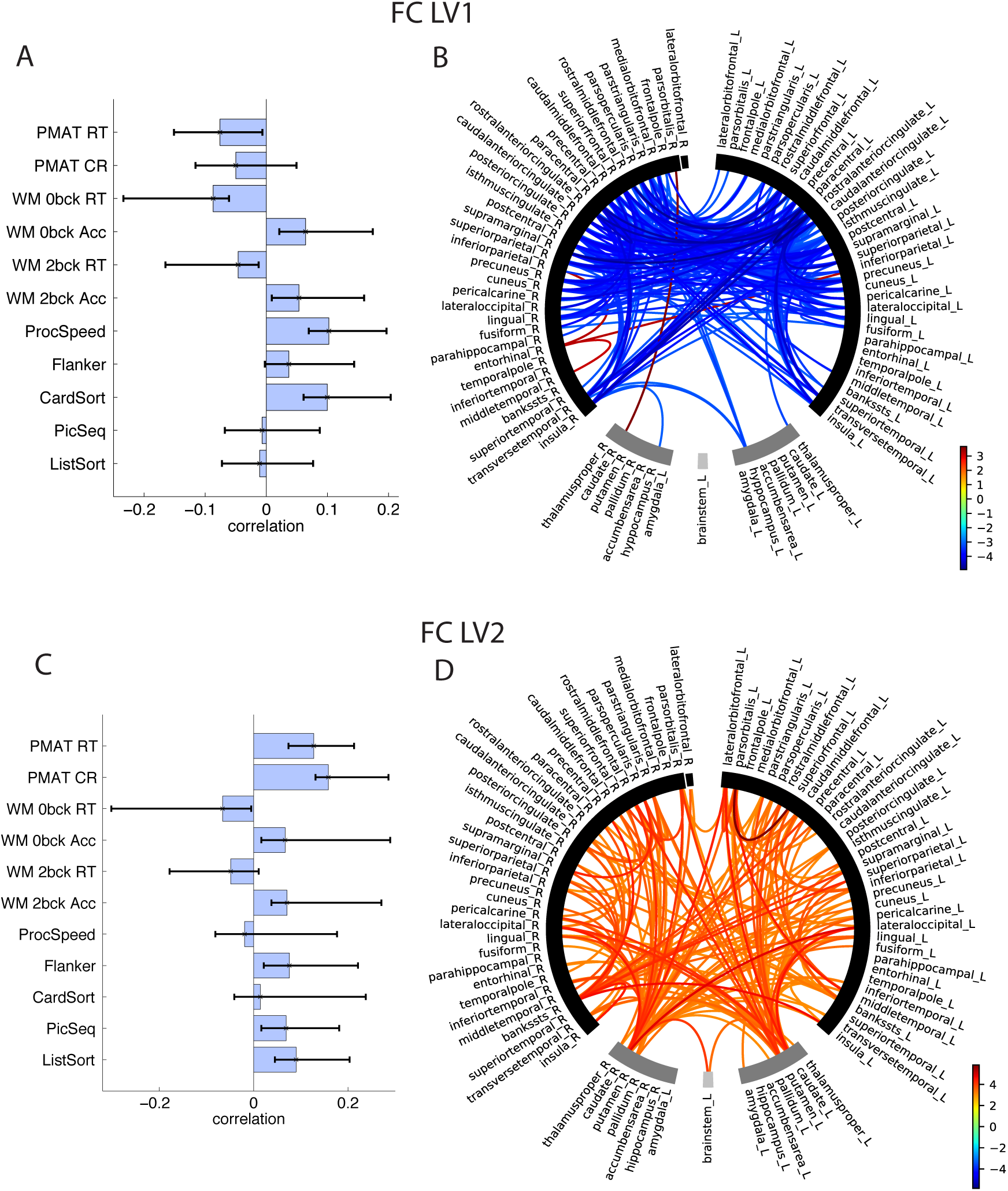
FC-Cognition A) LV1 correlations between FC and cognition, with CIs from bootstrap resampling, B) LV1 bootstrap ratios, these are connection loadings on the FC-Cognition relationship, bootstrap threshold = -3, 3. Connections with high bootstrap ratios contribute positively to the FC-cognition correlation, low bootstrap ratios contribute negatively to the FC-cognition correlation. Right and left hemisphere regions, and subcortical and cortical regions, are separated by a space in the circular plot. B) LV2 correlations between FC and cognition D) LV2 bootstrap ratios.

The first LV revealed FC-cognition correlations expressed across an array of cognitive tests, and a large number of cortical, mainly fronto-parietal connections (as well as insula, transverse temporal) that load negatively onto this relationship (See bootstrap ratios in Figure 4B, 4D). Thus, better performance on card-sorting, processing speed, WM accuracy, as well as better performance on the three RT measures (better performance = lower RT) was associated with lower FC. Only 4 connections loaded negatively onto this LV, 3 of which were connections of the R entorhinal. The second LV revealed FC-cognition correlations that were expressed primarily with the PMAT cognitive tests. A large number of inter-hemispheric cortico-cortical and cortico-subcortical connections loaded positively onto this LV. Thus, better performance on the PMAT tasks correlated with higher FC.

#### SC-Cognition vs FC-Cognition

As evident from Fig 3 and 4, there were more connections in the FC that correlate with cognition compared to in the SC. This can be shown as the number of connections that exceeded the bootstrap ratio threshold for each connection-cognition LV, expressed as a proportion of the number of connections that entered the PLS analysis (3403 vs 870, FC, SC). The proportion of connections that exceeded the bootstrap threshold were as follows, FC: LV1 = 15%, LV2 = 9%; SC: LV1 = 2%, LV2 = 1%. We hypothesized that this may be due to the difference in total between-subjects variance in the SC and FC. To this end, we conducted a SVD of the subject-wise SC (609 subjects * 870 connections) as well as the subject-wise FC (609 subjects * 3403 connections) and compared the sum of the squared singular values corrected by the number of connections that entered the analysis, in order to account for the sparsity of the SC (ΣS^2^_corrected_). Indeed, we found that the total corrected variance in subject-wise SC was smaller than in the subject-wise FC (ΣS^2^_SC_corrected_ = 0.0198, ΣS^2^_FC_corrected_ = 0.0294).

We observed that a very different pattern of connections in the SC related to cognition than in the FC. This can be expressed quantitatively as the dot product of the PLS brain scores (“U”) from the FC-cognition SVD and from the SC-cognition SVD, across both LVs. These dot products were close to zero (Table 2). This comparison was made for the behaviour contributions to each LV as well. To this end, we calculated the dot product of the PLS cognitive scores (“V”) from the FC-cognition SVD and the cognitive scores from the SC-cognition SVD (Table 3). In order to map which cognitive PCs were expressed in which LVs, we correlated the cognitive PC loadings with the SC-cognition and FC-cognition correlations on each LV (Table 4). Note that these correlations were all significant, *p* < 0.001, MC corrected (See Methods). We found that some cognitive PCs were particularly strongly captured in specific connectivity-cognition LVs. For example, the first cognitive PC was expressed very strongly in the first SC-cognition LV. Yet, even the second SC-cognition LV correlated well with this PC.

**Table 2.**
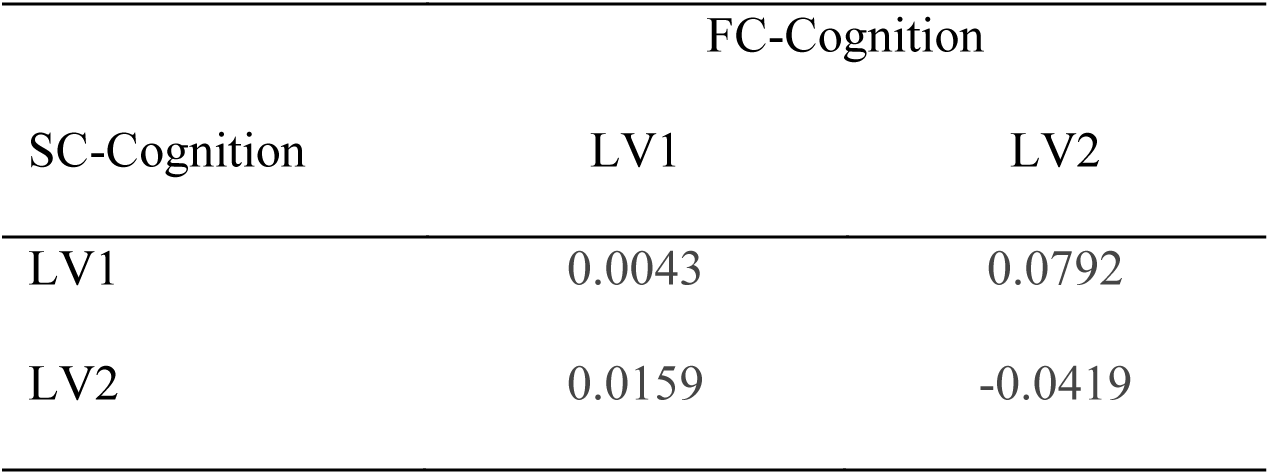
Dot product between brain scores for FC-cognition and SC-cognition across both LVs.

**Table 3.** Dot product between behavioural scores for FC-cognition and SC-cognition across both LVs.

| SC-Cognition | FC-Cognition |  |
| --- | --- | --- |
|  | LV1 | LV2 |
| LV1 | 0.51 | 0.83 |
| LV2 | 0.38 | -0.07 |

**Table 4.** Correlations of cognitive PCs loadings and SC-Cognition and FC-Cognition

| Cognition | SC-Cog |  | FC-Cog |  |
| --- | --- | --- | --- | --- |
|  | LV1 | LV2 | LV1 | LV2 |
| PC1 | 0.9825 | 0.7415 | 0.6798 | 0.6473 |
| PC2 | -0.2102 | -0.4327 | -0.8290 | 0.3725 |
| PC3 | -0.5635 | -0.2377 | -0.0664 | -0.5344 |

## Discussion

In the present study, we compared whole-brain cortical and subcortical SC and rsFC from 609 subjects from the Human Connectome Project with individual cognitive function across an array of 11 measures, including measures of working memory (WM n-back accuracy and RT, and ListSort), executive function/cognitive flexibility (CardSort), processing speed (ProcSpeed), fluid intelligence (PMAT), episodic memory (PicSeq), and attention/inhibitory control (Flanker). We first showed that the cognitive measures mapped onto three principal components (PCs) reflecting the heterogenous nature of functions measured via the tests. PC1 was a global cognition component that emphasized a speed-accuracy trade-off for the two WM tests. PC2 expressed fluid intelligence (PMAT). PC3 emphasized a more deliberate, slower, executive function/cognitive flexibility and attention/inhibitory control.

The conclusions of the study are two-fold: 1) that unique sets of connections map onto specific components of cognitive function, and 2) that SC and FC each capture independent and unique connections that relate to cognition. Importantly, previous approaches to studying connectivity-cognition relations have constrained the dimensionality via PCA before conducting Canonical Correlation Analysis (CCA) (Smith, 2016), which constrains how the behaviour can be projected into the brain space. In contrast, we compared the conjunction of brain and behaviour without first constraining dimensionality by PCA. Moreover, unlike CCA, PLS is robust to the colinearity that may be present in connectivity data. Thus, our method may be more sensitive in being able to extract the dimensions that relate brain and behaviour.

### Unique sets of connections map onto specific components of cognitive functions

First, we showed that SC and rsFC each capture multiple sets of connection-cognition associations. Two LVs for SC, and two for FC, each expressed a distinct and unique set of connections and components of cognitive function. This is in line with the view that distinct sets of connections are required to support heterogenous components of cognitive function, implying more specialized connectivity (Rosenberg et al., 2013) as opposed to the view that a global cognitive factor can be captured via a single set of connections (Malpas et al., 2016; Smith, 2016).

The two SC-cognition LVs each captured a limited, unique set of primarily intra-hemispheric cortical and subcortical connections. The first SC-cognition LV mapped almost perfectly onto the first global cognition PC, which can be attributed to the low variability across SCs. The second SC-cognition LV strongly expressed fluid intelligence (PMAT test) via a set of distributed cortical and subcortical connections.

The two FC-cognition LVs each captured a large set of unique inter and intra-hemispheric connections. The connectivity-cognition correlations we found were comparable to those identified previously with the HCP dataset (i.e., *r* = 0.045) (Hearne et al., 2016). The first FC-cognition LV expressed fronto-parietal, visual, and cingulate cortical connections, as well as transverse temporal, insula and L hippocampus connections to the rest of the cortex. rsFC in these regions was negatively associated with processing speed, executive function/cognitive flexibility, and working memory performance. The second FC-cognition LV was expressed via widely distributed cortico-cortical and cortico-subcortical connections, with strong contributions from the bilateral caudate and putamen connections to cortical regions. A large number of inter-hemispheric connections were expressed in this LV. The PMAT fluid intelligence measures were particularly dominant, with higher rsFC correlating with higher PMAT correct response as well as reaction time scores, so that subjects with higher rsFC were more accurate, but slower. We note that for both SC and FC, the PMAT measures were strongly expressed in the second LVs. The PMAT is an abbreviated version of the Raven’s Progressive Matrices (Prabhakaran et al., 1997; Gray et al., 2003; Wendelken et al., 2008), and is a measure of fluid intelligence that has been previously linked to individual differences in FC (Finn et al., 2015; Hearne et al., 2016; Smith, 2016; Ferguson MA, 2017).

The patterns of connectivity that were correlated with cognitive variability in our study were consistent with those observed previously. On the side of SC, the limited, widespread connections that associated with cognition were consistent with what had previously been reported by Ponsoda, Martinez et al (2017), where only 36 connections distributed across the entire brain predicted higher order cognitive functions. A number of these connections mapped closely onto the SC connections that we identified across the two LVs. On the side of the rsFC, connectivity distributed across the cortex likely supports cognitive function (Ferguson MA, 2017). Positive rsFC-cognition associations have typically been identified in fronto-parietal regions (Hearne et al., 2016), and negative associations in the default mode and dorsal attention network including visual and cingulate-parietal connectivity (Song et al., 2008; Song et al., 2009; Pamplona et al., 2015; Santarnecchi et al., 2015). Our first and second FC-cognition LV captured these negative and positive associations, respectively. Our second FC-cognition LV expressed several cortico-subcortical connections, primarily between the putamen, caudate, thalamus and the rest of the cortex. This is not surprising, as the cortical control of behaviour is mediated via several cortico-striatal-thalamic-cortical circuits (Alexander et al., 1986; Peters et al., 2016) that can be identified via structural and functional imaging (Seeley et al., 2007; Metzger et al., 2010). Connectivity from subcortical areas including the striatum have previously been tied to individual differences in phenotype and behaviour (Vaidya and Gordon, 2013). The cortico-striatal-thalamic cognitive control loop also includes the brainstem (Peters et al., 2016), which exhibited cognition-related connectivity with the caudate in our LV2. Moreover, the striatum (putamen and caudate) is specifically involved in learning, storing and processing memories (Packard and Knowlton, 2002), and higher WM performance has been tied to higher connectivity between the cingulo-opercular network and putamen rsFC (Tu et al., 2012). In our first LV, the only subcortical region that had negative rsFC associations with cognition was the L hippocampus, consistent with prior work (Salami et al., 2014). Insular connectivity, primarily with anterior cingulate regions, was also particularly prominent in the first LV, a region that may be important for reactive attentional control (Jiang et al., 2015).

Interestingly, in a sample of 317 HCP subjects, Hearne et al. (2016) identified only positive network-level associations between rsFC and two cognitive measures (one of these was the same PMAT measure used in the present study). As the authors themselves note, this may be because their network-level approach overlooks any existing edge level connectivity-cognition relationships that may exist (Song et al., 2008; Song et al., 2009; Pamplona et al., 2015; Santarnecchi et al., 2015). We replicated the positive association between rsFC and PMAT scores that was found previously (Hearne et al., 2016) within our second FC-cognition LV, where fluid intelligence was emphasized as the strongest cognitive correlate.

### SC and FC each capture independent and unique connections that relate to cognition

Second, we showed that that SC and FC each captured independent and complementary features of the connectome that were linked to cognitive function. The connections that expressed the SC-cognition association did not overlap with behaviourally-relevant connections in the FC, evidenced by the comparison of the spatial pattern of brain scores between the two analyses. While a similar suggestion has previously been made for SC versus task FC (Duda et al., 2010) and rsFC in select pathways (Hirsiger et al., 2016), it has not as of yet been examined in whole-brain SC-rsFC in as large a sample as ours. We found that far fewer connections within the SC compared to the FC associated with cognition, even when correcting for the sparsity of the SC. This was likely due to the comparably smaller amount of total variance in the SC across subjects. The limited, yet distributed nature of the SC network that varies with cognition has previously been reported (Ponsoda et al., 2017). Yet, we note that the amount of covariance accounted for by SC and by FC towards cognition was comparable. This suggests that both modalities are equally important for understanding individual differences in cognition.

It is important to consider that the imperfect association between SC and FC (Koch et al., 2002; Skudlarski et al., 2008; Honey et al., 2009) may impose limitations on the amount of overlapping information that can be provided by the two modalities. Moreover, the large repertoire of cognitive brain function is made possible by virtue of the dynamic nature of FC, which arises from a static SC backbone (Park and Friston, 2013). In this vein, an investigation into functional connectivity dynamics may help describe how the spatial contributions of SC and rsFC to cognition fluctuate over time. This investigation is, however, beyond the scope of this study.

## Acknowledgments

Data were provided by the Human Connectome Project, WU-Minn Consortium (Principal Investigators: David Van Essen and Kamil Ugurbil; 1U54MH091657) funded by the 16 NIH Institutes and Centers that support the NIH Blueprint for Neuroscience Research; and by the McDonnell Center for Systems Neuroscience at Washington University. The authors acknowledge the support of the NSERC grant (RGPIN-2017-06793). The authors declare no competing financial interests.

